# Secreted folate receptor-gamma drives fibrogenesis in nonalcoholic steatohepatitis by amplifying TGFβ signaling in hepatic stellate cells

**DOI:** 10.1101/2022.07.21.500829

**Authors:** Connor R. Quinn, Mario C. Rico, Carmen Merali, Oscar Perez-Leal, Victoria Mischley, John Karanicolas, Scott L. Friedman, Salim Merali

## Abstract

Hepatic fibrosis is the primary determinant of mortality in nonalcoholic steatohepatitis (NASH) patients. Antagonism of transforming growth factor β (TGFβ), a master profibrogenic cytokine, is a promising therapeutic target that has not yet been translated into an effective therapy, due in part to the lack of animal models resembling the human phenotype of NASH. Here we have identified that soluble secreted folate receptor gamma (FOLR3), expressed in humans but not rodents, is a secreted protein that is elevated in livers of NASH subjects but not in subjects with nonalcoholic fatty liver, type II diabetes, or healthy subjects. FOLR3, based on global proteomics, was the most highly expressed NASH-specific protein and positively correlated with increasing fibrosis stages, suggesting an impact on activated hepatic stellate cells (HSCs), the key fibrogenic cell in the liver. Exposure of stellate cells to exogenous FOLR3 led to elevated extracellular matrix (ECM) protein production, an effect synergistic with TGFβ1. Structurally, FOLR3 interacts with serine protease HTRA1, which downregulates TGFβ signaling through the degradation of its receptor TGFBR2. Administration of human FOLR3 to mice induced severe bridging fibrosis and an ECM pattern resembling human NASH. Our study uncovers a novel role of FOLR3 in enhancing fibrosis and identifies FOLR3 as a potential therapeutic target in NASH fibrosis.

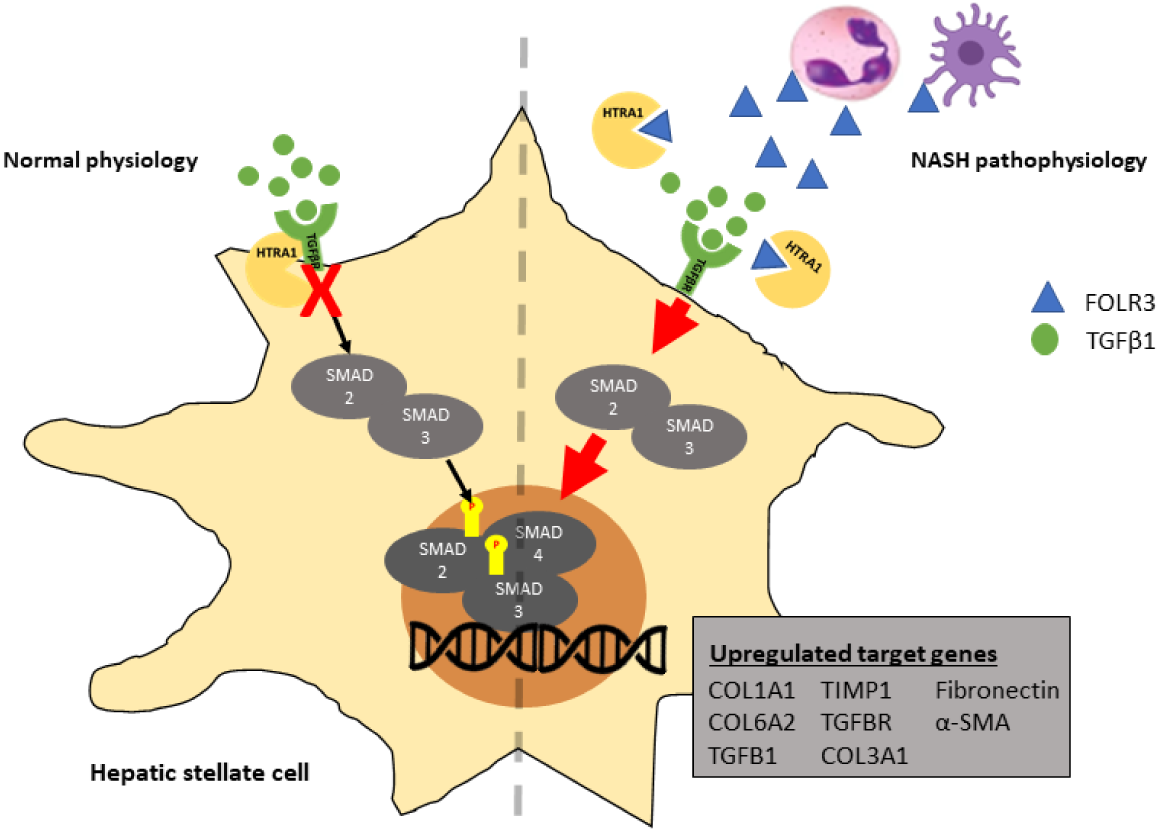

## Introduction

Obesity has rapidly become a worldwide epidemic, magnifying the effects of the metabolic syndrome^1-3^. Nonalcoholic fatty liver disease (NAFLD) is the hepatic manifestation of metabolic syndrome and can negatively affect a patient’s well-being. NAFLD is a spectrum of diseases ranging from simple steatosis to steatohepatitis, eventually cirrhosis, and complete liver failure. The prevalence of nonalcoholic steatohepatitis (NASH) has risen dramatically in the past ten years and significantly increases the risk of decompensated liver failure and liver transplant^4^. NASH is characterized by steatosis, inflammation, and varying degrees of fibrosis. Previous studies have demonstrated that fibrosis progression is the number one determinant of adverse events in NASH patients, and fibrosis resolution is key for treating NASH^5-7^. Despite the growing impact of NASH on the healthcare system and the importance of fibrosis, there is still an absence of approved therapies for treating NASH and liver fibrosis. Thus, there is a significant unmet need for understanding liver fibrogenesis and identifying effective drug targets for treating NASH.

NASH arises through multiple metabolic insults, including obesity, insulin resistance, and lipotoxicity, leading to inflammation, hepatocellular death, and fibrosis^8-10^. Although a complete understanding of the molecular mechanisms causing these insults is still lacking, several studies have begun demonstrating the importance of hepatocytes and non-parenchymal cell types in driving the disease^11-13^. Liver fibrosis involves the accumulation of extracellular matrix proteins (ECM) in response to chronic liver injury. Hepatic stellate cells (HSCs) have been identified as the primary mediators of extracellular matrix protein production and secretion during fibrosis^14^. Under normal conditions, HSCs remain in a quiescent state responsible for storing vitamin A. In response to liver injury and inflammation, HSCs transdifferentiate into a myofibroblast phenotype expressing high levels of ECM^15^. Transforming growth factor beta 1 (TGFβ1) has been identified as the most potent fibrogenic cytokine by activating transcription factors SMAD2/3. A better understanding of stellate cell fibrogenesis during NASH fibrosis could lead to mechanism-based therapies for blocking fibrosis progression and improving patient outcomes.

FOLR3 is part of a specialized folate receptor family with a conserved binding region for folate metabolites^16^. These receptors can mediate the intracellular uptake of folate, but their role in folate metabolism and overall function is still not well understood. FOLR3 differs from the other folate receptors by lacking a glycophosphatidylinositol (GPI) modification signal. Modification by GPI acts as a hydrophobic anchor to attach the receptor to the cell membrane and mediate folate intake into the cell. Interestingly, FOLR3 is not attached to the cell membrane and is a secreted protein with a similar affinity for folates as FOLR1 and FOLR2 ^17^. In addition, FOLR3 is mainly expressed in hematopoietic tissues and is present in the systemic circulation^17,18^. Since the discovery of FOLR3, there have been very few reports clarifying its function ^18,19^.

In this study, we performed a global unbiased proteomic analysis that identifies increased expression of FOLR3 in the livers of NASH subjects. We then characterized its impact on ECM remodeling using in vitro assays in HSCs and protein-protein interaction studies. Finally, we determined the role of FOLR3 in exacerbation of fibrosis in a diet-induced mouse model of NASH. We successfully applied these studies to uncover a novel role of FOLR3 in human hepatic fibrogenesis.

## Results

### Hepatic proteome profiling identifies FOLR3 in NASH

NASH is commonly associated with other metabolic conditions, including obesity, type II diabetes, and steatosis. An unbiased global proteome pipeline has not been extensively applied to human NASH livers with appropriate controls, and therefore we first set up to study phenotypical proteome changes^20-22^. Since NASH is a more severe disease than steatosis, we sought to identify proteins/pathways critical to NASH. We interrogated a cohort of liver tissue samples from four groups of subjects (n=6 in each group): NASH, steatosis, type II diabetes (T2DM), and healthy (Table 1. and Extended Data Table 1). Liver tissue samples were prepared and analyzed using a label-free global proteomic analysis workflow as previously described^23-25^ (Extended Data Figure 1). Among the study groups, 5,860 proteins were identified and quantified with high confidence (Extended Data Table 2). The grouped heat map shown in Figure 1 displays the differentially expressed proteins among the cohorts, and unsupervised clustering demonstrates that NASH has the most distinct proteomic profile. Several proteins are differentially expressed in the NASH liver that has previously been associated with NASH, including proteins regulating liver metabolism, ENO3, and Hp, and inflammation/apoptosis, SCAF11, FAM169A, and GSTM5^26-30^. Of the differentially expressed proteins, we identified folate receptor gamma (FOLR3) as the highest upregulated protein that is specific to the NASH cohort. Further proteomic analysis of a cohort of NASH patients with varying fibrosis stages showed that FOLR3 levels significantly correlate with fibrosis stage (Figure 1).

**Table 1.** Subject characteristics for global proteomic analysis

|  | Healthy (n=6) | NASH (n=6) | Steatosis (n=6) | T2DM (n=6) |
| --- | --- | --- | --- | --- |
| Age | 40.5 ± 17.0 | 57.2 ± 15.9 | 53.7 ± 6.3 | 48.0 ± 9.5 |
| Gender: M, F (%) | 50,50 | 33.3,66.7 | 50,50 | 66.7,33.3 |
| BMI | 20.0 ± 3.15 | 39.9 ± 14.9 | 37.5 ± 4.6 | 37.9 ± 7.5 |
| Type II Diabetes (%) | 0 | 0 | 100 | 100 |

**Figure 1.**
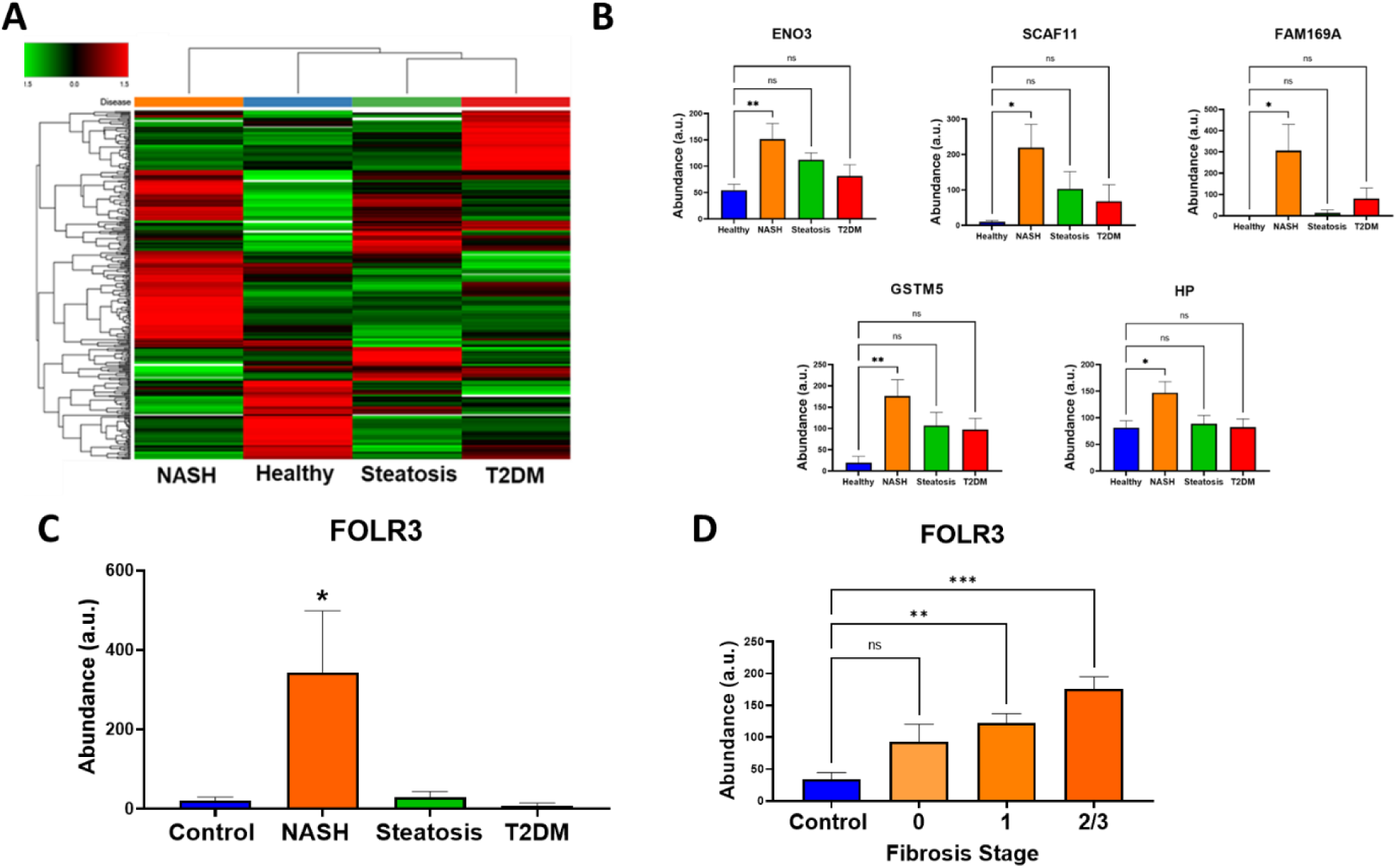
Folate receptor gamma is increased in NASH liver. (A) Heat map of differentially expressed proteins among cohorts with (B) quantitation for selected proteins. (C) Individual FOLR3 abundance from the global proteomic analysis. (D) Correlation analysis between FOLR3 abundance and fibrosis stage. (n=6 * p>0.05 ** p<0.01 *** p<0.001)

To rigorously quantify FOLR3, a targeted mass spectrometry method using nano-liquid chromatography and selected reaction monitoring (SRM) was performed. SRM is a highly sensitive and specific label-free mass spectrometry method based on quantitating a unique tryptic peptide belonging to the protein of interest ^31^. The SRM method development, instrument parameters, and ion transitions used for quantitation and standard curve are shown in Figure 2 and Extended Data Figure 2 and Table 3. The accuracy and precision of the method were assessed using six (n=6) replicates of mouse liver lysate spiked with two different concentrations of FOLR3, 50pg/ug of protein (low QC), and 1000pg/ug of protein (high QC). The accuracy and precision of the high QC were 18% and 13.1%, and the low QC was 17.6% and 9.1%, respectively. The method was used to quantify FOLR3 in liver samples from subjects with NASH, T2DM, steatosis, or healthy controls. The results in Figure 2 show that FOLR3 is markedly increased in NASH, correlating with our findings in the global proteomic analysis. There was a 10-fold increase in FOLR3 levels in control versus NASH subjects with an average FOLR3 concentration of 184 ± 200 pg/ug protein and 1826 ± 560.7 pg/ug of protein for controls and NASH, respectively.

**Figure 2.**
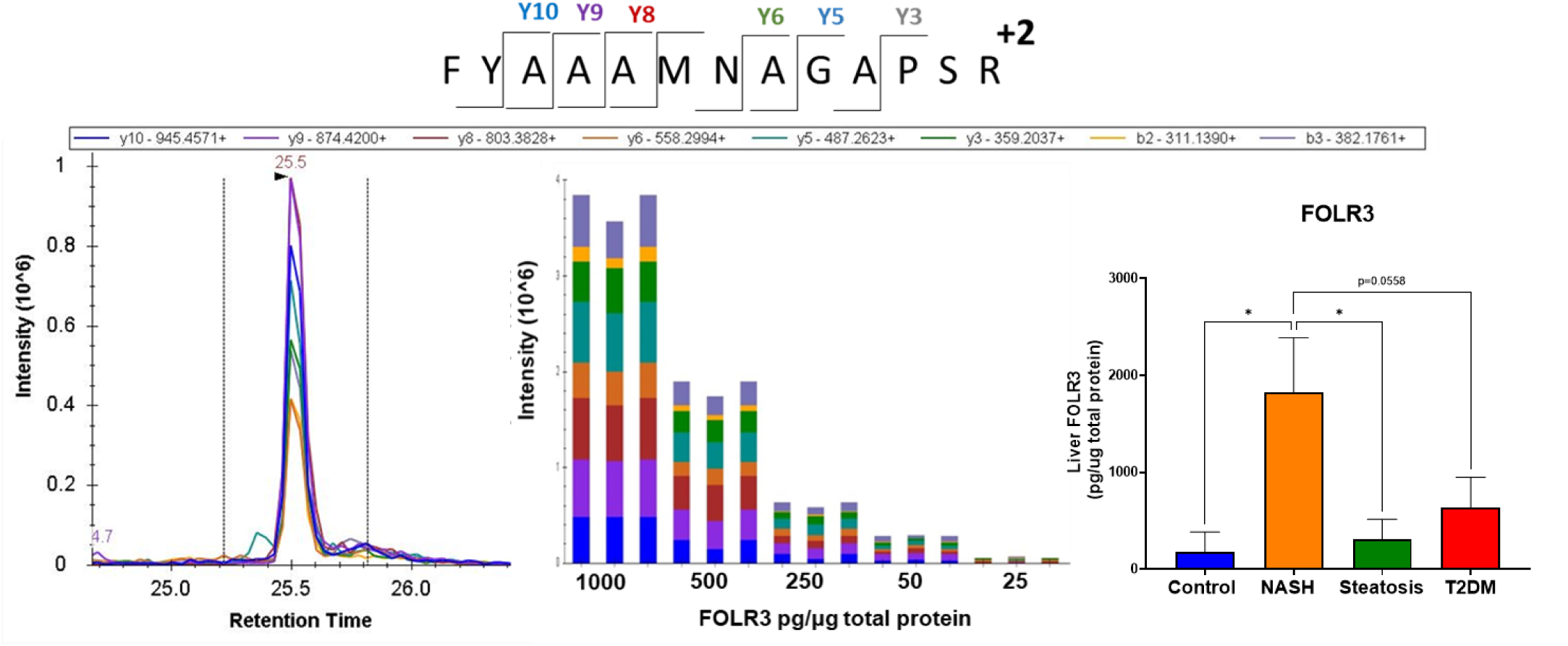
FOLR3 quantitation using targeted mass spectrometry. A selected reaction monitoring method was developed for a unique tryptic peptide of FOLR3. The fragmentation of the precursor ion, the coelution of the product ions, the peak area response to concentration standards, and quantitation of FOLR3 in human liver samples using selected reaction monitoring method are shown. (n=5 *p<0.05)

### FOLR3 enhances TGFβ1-mediated HSC activation

FOLR3 has not yet been characterized in hepatic metabolism or fibrosis, prompting further studies into its function using immortalized primary human HSCs (LX-2). To assess the activation of HSCs by FOLR3 itself, we used global proteomic analysis to analyze the secretome of HSCs treated with FOLR3 and TGFβ1 in serum-free media^25^. Cells were treated with vehicle, FOLR3 1 ug/ml, TFGβ1 2 ng/ml, or FOLR3 1 ug/mL and TGFβ1 2 ng/mL for 24 hours (n=4). The FOLR3 concentration was chosen based on its ability to increase the production of COL1A1, a common marker for HSC activation, in a concentration-dependent manner (Extended Data Figure 3). The cell supernatants were individually processed and prepared for global unbiased label-free proteomic analysis as described in the methods. We detected and quantified 1,707 proteins from this analysis, with 97 extracellular matrix proteins (Extended Data Table 4). The heat map displayed in Figure 3 shows the expression of extracellular matrix proteins from the four experimental groups. Hierarchical clustering shows proteins specifically upregulated in cells treated with FOLR3: collagen alpha-1(XVIII) chain and collagen alpha-1(X) chain or TGFβ1: CCN family member 2, collagen alpha-2(IV), plasminogen activator inhibitor 1, and matrix metalloproteinase-3. The cells treated with FOLR3 and TGFβ1 demonstrated a synergistic effect increasing the expression of several extracellular matrix proteins, including collagen and matrix metalloproteinases. In addition, TGFβ1 and TGFβ regulatory proteins, latent transforming growth factor binding protein 2 and 3 (LTBP2 and LTBP3) and transforming growth factor beta-induced protein (TGFBI) were all significantly increased with the treatment of FOLR3 and TGFβ1, demonstrating FOLR3 can have a direct effect on TGFβ signaling and remodel the ECM (Figure 3). Principal component analysis (PCA) corroborates these findings showing the FOLR3+TGFβ1 replicates clustered together, demonstrating a distinct expression profile compared to the other experimental groups. The secretion of COL1A1 in cell culture media was further analyzed by western blotting (Figure 3C). These results corroborated the elevation of intracellular COL1A1 in the cells treated with FOLR3 and TGFβ1 (Extended Data Figure 3). Overall, these results demonstrate that FOLR3 increases extracellular matrix protein production in HSCs and enhances the effect of TGFβ1 on fibrogenesis.

**Figure 3.**
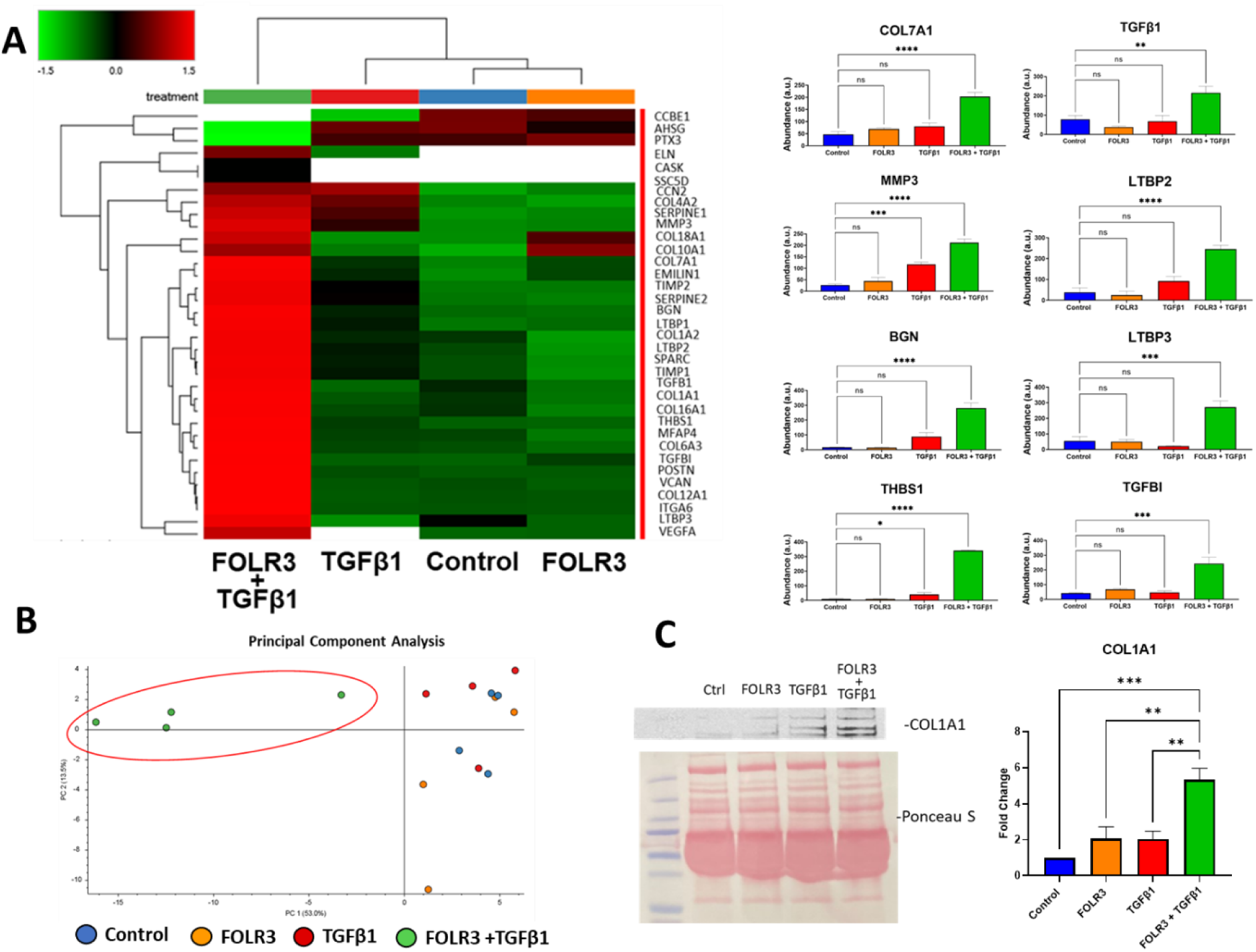
FOLR3 enhances TGFβ1 mediated HSC activation. (A) Heat map showing the expression of extracellular matrix proteins in the supernatant of cells treated with vehicle, FOLR3, TGF1β, and FOLR3+TGF1β and individual quantitation of NASH-associated proteins (n=4 *p<0.05 **p<0.01 ***p<0.001) (B) Principal component analysis plot (C) Western blot analysis and quantitation of COL1A1 (n=3 **p<0.01 ***p<0.001)

### FOLR3 regulates TGFβ signaling through interaction with serine protease HTRA1

To determine how FOLR3 activates HSCs’ ECM production, protein-protein interaction studies were performed. First, immunoprecipitation using magnetic beads coated with anti-FLAG antibodies were used to pull down FLAG tagged-FOLR3 with interacting proteins. FOLR3 and interacting proteins were then eluted from the magnetic beads and analyzed using LC-MS/MS. Samples with and without FOLR3 were processed simultaneously to eliminate false positives. We identified three proteins specifically pulled down by FOLR3, serine protease HTRA1, glutaminyl peptide cyclotransferase, and midkine (Extended Data Table 5). Interestingly, all these proteins are involved in ECM metabolism. The most abundant protein in the pulldown was serine protease HTRA1, which was only detected in the pulldown samples when FOLR3 was present (Figure 4A). The peptide spectrum match for the most intense HTRA1 peptide is shown in Figure 4B and provides high confidence for detecting HTRA1. HTRA1 is involved in extracellular matrix remodeling and regulates TGFβ signaling through degradation of transforming growth factor receptor 2 (TGFBR2)^32,33^. Graham et al. previously demonstrated HTRA1 can degrade TGFBR2 on the cell surface in a dose-dependent manner and silencing HTRA1 increases TGFβ responsiveness. To further confirm the interaction between FOLR3 and HTRA1, the pulldown experiment was repeated and analyzed using Western blot (Figure 4C). These results confirmed the interaction of FOLR3 and HTRA1.

**Figure 4.**
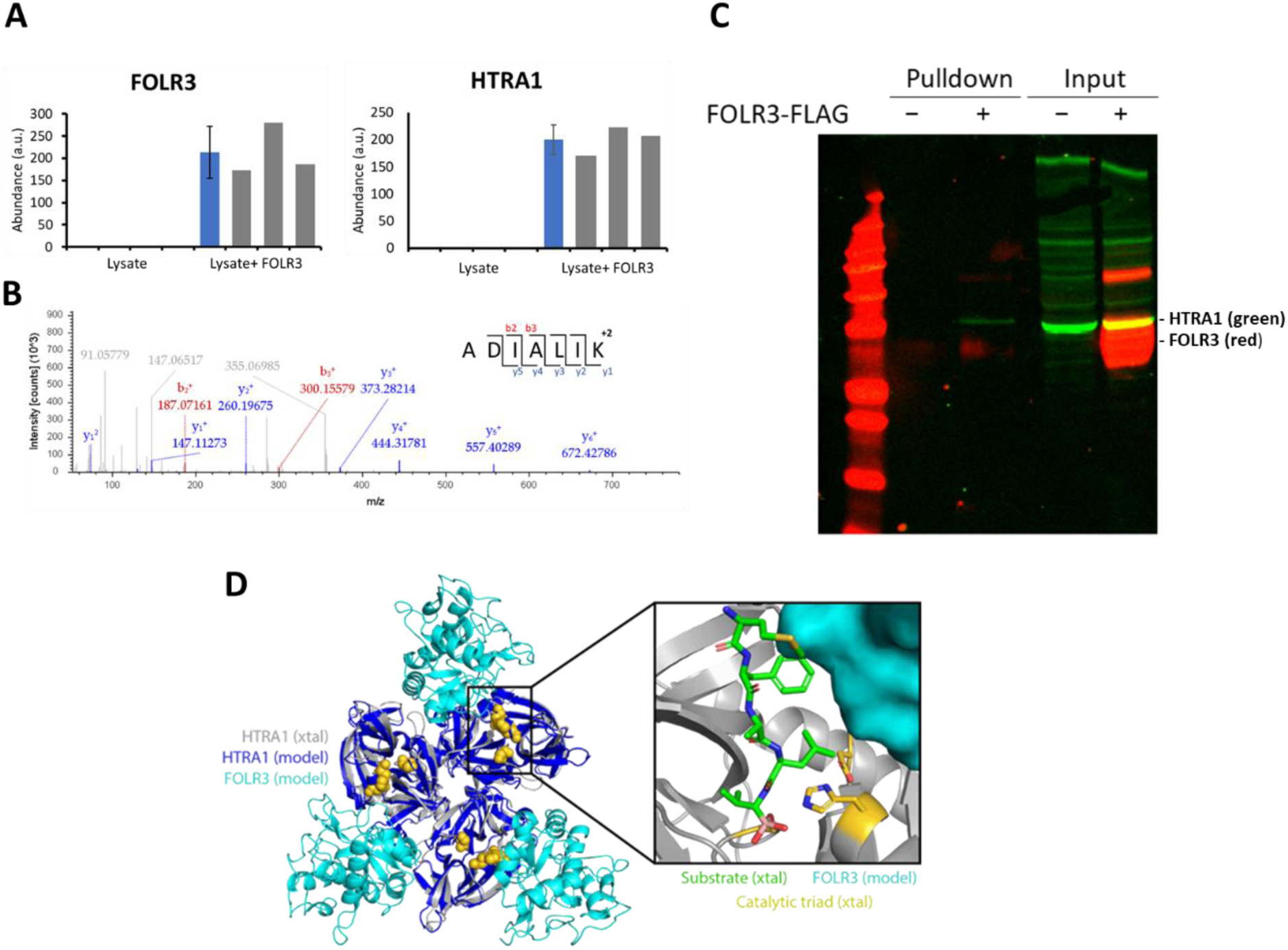
Folate receptor-gamma interacts with HTRA1. Anti-FLAG coated magnetic beads were used to pulldown FOLR3 and interacting proteins within HSC lysate. Samples without FOLR3 were processed simultaneously. All pulldowns were analyzed by LC-MS/MS. (A) Scaled abundance for FOLR3 and HTRA1 in pulldown samples. Blue bars are average abundance and standard deviation for each group, and gray bars are individual abundances for each replicate (B) Peptide spectrum match for serine protease HTRA1. (C) Western blot analysis for HTRA1 after FOLR3 pulldown. (D) AlphaFold2 model of the HTRA1-FOLR3 interaction. The trimeric arrangement of HTRA1 in the model (blue) is essentially identical to the previously-reported crystal structure of the HTRA1 catalytic domain (silver). Three copies of FOLR3 (cyan) are symmetrically arranged relative to HTRA1 in the model, close to the active site (gold). As shown in the inset, this model of FOLR3 (cyan) does not include direct contacts with the residues comprising the catalytic triad (gold); rather, it occludes the region that would otherwise be occupied by substrate (green), suggesting a means by which FOLR3 may directly inhibit the proteolytic activity of HTRA1.

To explore the potential structural basis for the interaction between FOLR3 and HTRA1, we built a model of this interaction using ColabFold-AlphaFold2^34,35^. The catalytic (serine hydrolase) domain of HTRA1 is known to form a homotrimer, with the N-and C-terminal domains of the protein dispensable for its activity^36^. Accordingly, we applied ColabFold-AlphaFold2’s multimer model^37^ using a 3:3 stoichiometry of FOLR3 and HTRA1’s catalytic domain. In the best-scoring model (Figure 4D) the overall structure and trimeric arrangement of HTRA1 are virtually superposable with the previously-reported crystal structure^36^ and the three copies of FOLR3 are placed in a symmetric arrangement relative to each subunit. Specifically, FOLR3 occupies a region just outside the active site, engaging the protein close to the catalytic triad. While there are no direct contacts with the residues comprising the catalytic triad, the model of FOLR3 overlaps with a peptide inhibitor included in the crystal structure (Figure 4D, inset): thus, we speculate that the binding of FOLR3 to this region might inhibit catalysis by preventing substrate access to the active site.

Next, we explored if HTRA1 increases the degradation of TGBR2 in HSCs to reduce fibrogenesis. Immunofluorescence was used to visualize active TGFBR2 on the cell membrane of HSCs (Figure 5). Activated HSCs showed robust expression of TGBR2 on the cell surface. Treating the cells with HTRA1 for 2 hours significantly decreased the amount of TGFBR2 on the cell surface, and pre-treatment with FOLR3 abrogated the degradation of TGFBR2. These results indicate that FOLR3 can inhibit the degradation of TGFBR2 and establish the functional impact of FOLR3 on HTRA1. Altogether, these studies demonstrate that FOLR3 interacts with HTRA1 to prevent the degradation of TGFBR2.

**Figure 5.**
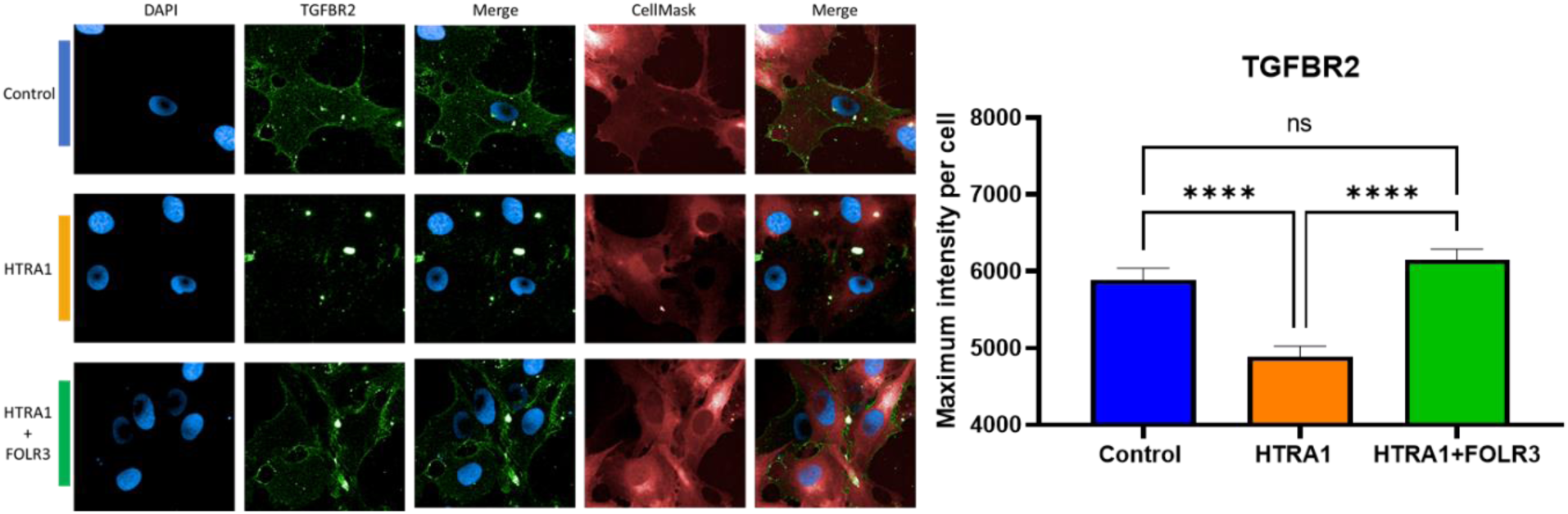
FOLR3 regulates TGFβ signaling through interaction with HTRA1. Immunofluorescence of TGFBR2 (green) in LX-2 cells treated with HTRA1 and FOLR3. Nuclei and cell body stained in blue and red, respectively. Quantitation of TGFBR2 is shown in the bar graph. (n=300 **** p<.0001)

### FOLR3 exacerbates fibrosis in a diet-induced mouse model of NASH

Our in vitro characterization in cultured HSCs demonstrated that FOLR3 enhances TGFβ signaling through interaction with HTRA1. Mice do not express FOLR3, and do not develop severe fibrosis and cirrhosis that can characterize human disease using murine models of NASH. Based on the high sequence similarity between human and mouse HTRA1, we predicted that FOLR3 could induce severe fibrosis in a NASH diet-induced model. To assess the effect of FOLR3 in an in vivo model of NASH, recombinant FOLR3 was administered to animals on an Amylin diet, which induces features of NASH^25^. Animals were fed a chow diet (n=3) or an Amylin diet (n=6) for the entirety of the study. For the last four weeks of the study, three Amylin diet mice were treated with FOLR3 1ug/day i.p. This dose was chosen because it was effective in producing ECM elevation in HSCs. Liver tissue was collected at the end of the study, and histology was analyzed using the NAFLD Activity Score (NAS). Animals treated with FOLR3 had significantly elevated steatosis, hepatic ballooning, and overall NAS score compared to Amylin animals. Sirius red staining was used to visualize collagen fibers within the liver tissue and quantify the fibrosis score. Animals treated with FOLR3 had significantly higher levels of fibrosis and presented clear signs of advanced fibrosis, including bridging fibrosis commonly seen in human NASH. To evaluate changes at the molecular level, we performed an unbiased proteomic analysis on the liver tissue of each mouse. Several ECMs significantly increased in both Amylin and Amylin+FOLR3 animals (Extended Data Table 6). Furthermore, the heat map shown in Figure 6 shows a subset of ECMs specifically increased in only Amylin+FOLR3 animals, including galectin-3 (Lgals3), apolipoprotein E (APOE), and proteins increased in our in vitro studies, thrombospondin 1 (THBS1), serpin family E member 2 (SERPINE2), and collagen type 6 (COL6A2). Interestingly, these proteins have previously been increased during TGFβ1 activation^15,38-40^. These results are in good agreement with our in vitro analyses and demonstrate that FOLR3 can drive NASH fibrogenesis in an animal model of NASH.

**Figure 6.**
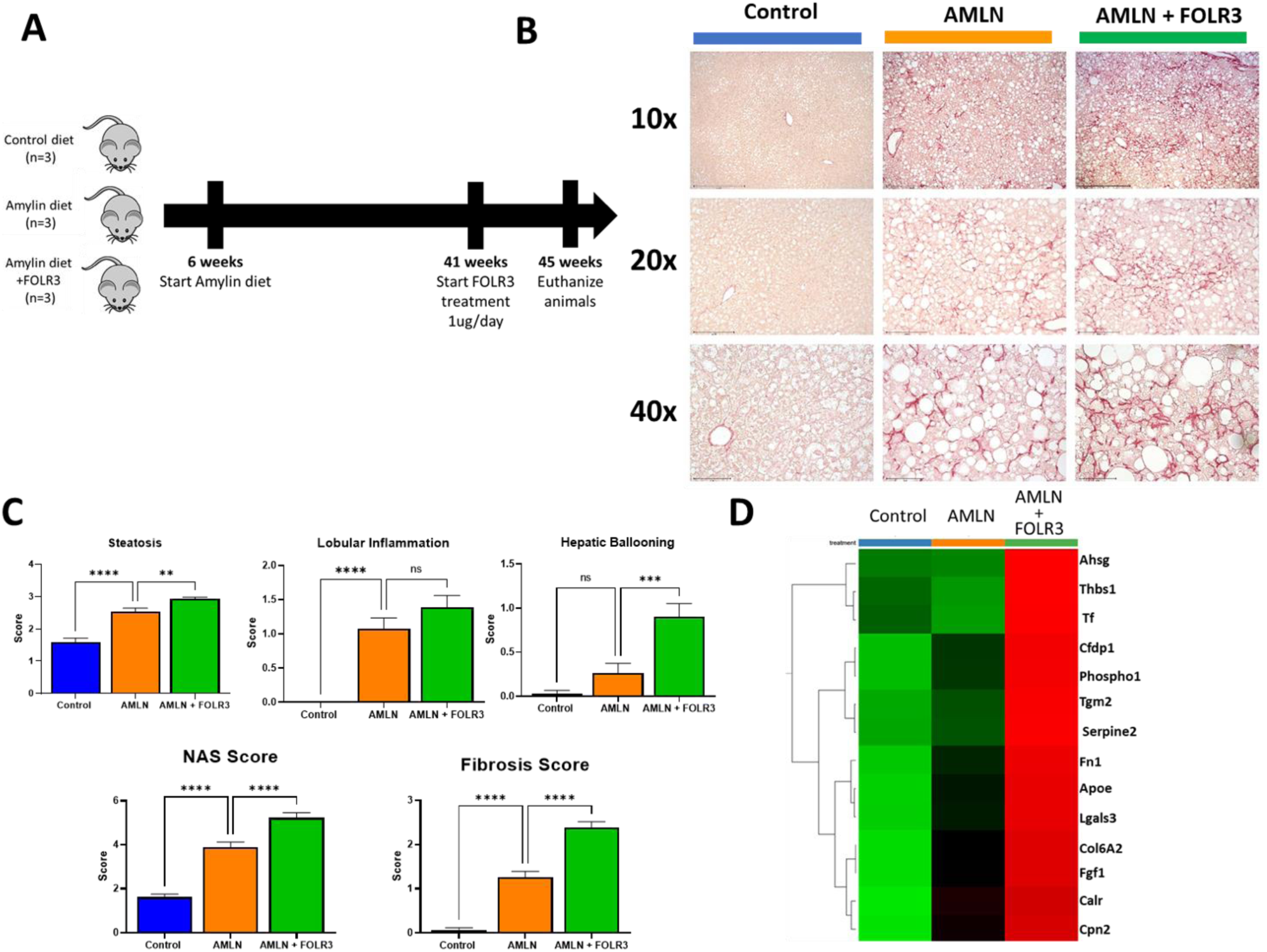
FOLR3 exacerbates fibrosis in a diet-induced mouse model of NASH. (A) Experimental design. (B) Mouse liver histology with Sirius red staining. Images are taken at 10x, 20x, and 40x. Scale bar = 500, 200, and 75µm, respectively (C) NAFLD Activity Score and Fibrosis score quantitation (D) Heat map displaying ECM abundance from proteomic analysis (n=3 **p<0.01 ***p<0.001)

## Discussion

FOLR3 is the secreted form of the folate receptor family and is present in the systemic circulation^18^. In this study, FOLR3 was identified as specifically upregulated in human NASH liver where it regulates TGFβ1 signaling through interaction with serine protease HTRA1. The folate receptor family, specifically FOLR1 and FOLR2, has been significantly overexpressed in various tumors and diseases associated with inflammation and targeted for therapeutic drug delivery^18,41,42^. Recent studies have identified that FOLR3 gene expression is increased in oral cancers and tumor peripheral blood mononuclear cells, but the molecular function of FOLR3 has not yet been characterized. The importance of FOLR3 has been demonstrated in studies reporting that FOLR3 single nucleotide polymorphisms (SNPs) are strongly associated with disease progression^19^, but the functional impact of these SNPs is not known, and not addressed in our study. Recent data have shown that folate receptors have additional roles independent of folate metabolism. For example, FOLR1 was shown to minimally impact folate uptake compared to reduced folate carrier (RFC) in multiple cancer cell lines ^43^. Furthermore, FOLR1 can act as a transcription factor. Hansen et al. reported that FOLR1 can activate pro-oncogene signal transducer and activator of transcription 3 (STAT3) and increase cancer cell proliferation^44^. Mayanil et al. described FOLR1 as a direct transcription factor in increasing the expression of several genes involved in pluripotency and stem cell regulation ^45-47^. These findings bring a novel interest in the role of folate receptors in disease pathophysiology.

This study identifies elevated FOLR3 expression in NASH for the first time, which has a positive correlation with fibrosis stages. Our proteomic analysis did not detect significant changes in other folate receptors or folate metabolic enzymes. Previous liver tissue single-cell RNAseq analyses showed that FOLR3 expression is predominantly in inflammatory macrophages^48^. Therefore, it is likely that FOLR3 is acting as a cytokine secreted from inflammatory macrophages during NASH and contributing to fibrogenesis through activation of HSCs.

Our in vitro studies using HSCs demonstrate that exogenous FOLR3 increases ECM protein production in HSCs in synergy with TGFβ1. Secretome analysis of conditioned HSC media showed upregulation of several extracellular matrix proteins in cells treated with FOLR3 and TGFβ1 compared to treatment with FOLR3 or TGFβ1 alone. These included several proteins involved in TGFβ1 signaling and regulation, including TGFβ1, TGFBI, and latent TGFβ binding proteins 1-3 (LTBP 1-3). TGFβ1 is tightly regulated in a spatial manner. It is secreted in an inactive state bound to latent TGFβ binding proteins in the extracellular matrix^49^. The localization of TGFβ1 in the ECM allows for rapid mobilization in response to cellular stimuli and activation of TGFβ1. LTBP1-3 is increased in NASH and hepatocellular carcinoma and reflects activation of TGFβ1 signaling^50,51^. In addition, previous studies showed that the knockout of LTBP1 in mice prevents hepatic fibrogenesis^52^. The overexpression of several ECMs, specifically TGFβ1 and LTBP1-3, demonstrates that FOLR3 can activate HSCs in the presence of TGFβ1 and can impact TGFβ1 regulation directly.

Extensive protein-protein interaction studies were performed to identify binding partners of FOLR3. IP-MS analysis identified the interaction between FOLR3 and serine protease HTRA1. HTRA1 is a secreted protein with a highly conserved trypsin-like serine protease domain that is implicated in several pathologies involving aberrant extracellular matrix deposition, including fibrotic cirrhosis^53^. Previous studies have shown HTRA1 can negatively regulate TGFβ signaling through degradation of TGFBR2^32,33^. For example, missense mutations of the HTRA1 gene have been identified in cerebral autosomal recessive arteriopathy with subcortical infarcts and leukoencephalopathy (CARSIL). These mutations lead to decreased proteolytic activity, disinhibition of TGFβ1 signaling, and increased production of ECMs ^54^. These findings show HTRA1 is a crucial regulator of TGFβ1 signaling and could be involved in HSC activation during hepatic fibrogenesis. Using molecular modeling, we found that FOLR3 binds HTRA1 near the HTRA1 active site and can prevent substrate binding. Our functional analysis demonstrates that HSCs treated with HTRA1 degrade active TGFBR2 on the cell membrane. To our knowledge, this is the first time HTRA1 has been shown to degrade TGFBR2 in HSCs. Furthermore, we showed that FOLR3 inhibits the degradation of TGFBR2 and preserves active receptors on the cell surface, correlating with our molecular model. These results indicate that FOLR3 prevents negative regulation of TGFβ signaling by HTRA1 and leads to excessive production and secretion of ECMs.

A significant limitation for developing therapeutics for NASH is the lack of preclinical animal models that appropriately model human disease. Diet-induced rodent models consistently develop steatosis and inflammation but do not develop the extent of fibrosis commonly seen in the human disease. Rodents do not express the FOLR3 gene, and we hypothesized that the lack of FOLR3 in rodents prevents severe fibrosis progression. In addition, mice express HTRA1 with high sequence homology with the human protein, implying human FOLR3 can interact with mouse HTRA1 and prevent its activity, as shown in our in vitro studies. To test our hypothesis, we fed mice a NASH-inducing diet and treated them with human FOLR3 to assess the effects on fibrosis development. Animals treated with FOLR3 developed advanced fibrosis stages, including excessive collagen deposition and bridging fibrosis, a key hallmark of human NASH.

Furthermore, proteomic analysis of the liver tissue showed mice treated with FOLR3 had increased expression of a unique subset of ECMs compared to animals on the Amylin diet, including collagen 6A2, galectin-3, and SERPINE2. These proteins were also increased during our in vitro studies with FOLR3 and have previously been increased in human NASH^38,39,55,56^. Together these results demonstrate that FOLR3 treatment can induce NASH with advanced fibrosis stages in mice. Therefore, FOLR3 treatment could be used as a model for generating a human fibrosis phenotype in mice to accelerate preclinical development of therapeutics for hepatic fibrosis in NASH.

Overall, this study identifies a novel role of FOLR3 in human NASH. We have shown that the increased abundance of FOLR3 during NASH can enhance TGFβ1 signaling in HSCs by binding to HTRA1 and preventing the degradation of TGFBR2. This leads to the overactivation of TGFβ signaling and the excessive production of ECMs. These results demonstrate that FOLR3 can be a potential drug target for treating hepatic fibrosis, and FOLR3 treatment can be used in preclinical models to induce advanced stages of fibrosis that better represent the human disease.

## Methods

### Human Liver samples

Human liver tissue was purchased commercially from XenoTech. Liver tissue was supplied as frozen pre-lysate in the buffer. Samples were fully characterized with pathological diagnoses of NASH, steatosis, or type II diabetes.

### Global mass spectrometry analysis

Global proteomic analysis was performed on liver tissue as previously described ^25^ Mass spectra processing was performed with Proteome Discoverer v2.5. The generated de-isotoped peak list was submitted to an in-house Mascot server 2.2.07 for searching against the Swiss-Prot database (Release 2013_01, version 56.6, 538,849 sequences), Sequest HT database, and MS Amanda. All search parameters were set as follows: species, homo sapiens; enzyme, trypsin with maximal two missed cleavage; fixed modification, cysteine carboxymethylation; 10 ppm mass tolerance for precursor peptide ions; 0.02 Da tolerance for MS/MS fragment ions.

### Targeted mass spectrometry analysis

Liver tissue samples were prepared as described in the global mass spectrometry analysis. Protein quantification was performed on a TSQ Quantum Ultra triple quadrupole mass spectrometer (Thermo Scientific, Waltham, MA, USA) equipped with an Ultimate 3000 RSLCnano system with autosampler (Thermo Scientific, Waltham, MA, USA). The mobile phase consisted of Buffer A, 0.1% formic acid, and Buffer B, 85% acetonitrile with 0.1% formic acid. Peptides were separated on an Acclaim PepMap RSLC C18 precolumn (3 µm, 100 Å) and column (2 µm, 100 Å). They were then eluted with a 2–95% Buffer B flow gradient over 20 min. Peptides entered the mass spectrometer through nanoelectrospray ionization with a voltage of 1600 V and capillary temperature of 270 °C. The mass spectrometer was run in selected reaction monitoring mode detecting the transitions listed in Extended Data Table 3. Data was collected using Xcalibur software 4.1 and imported into Skyline software 21.1 for peak area integration and data analysis. FOLR3 peak area was normalized by β-actin for each sample.

### Cell culture

LX-2 cells were purchased from Sigma and grown in Dulbecco’s minimal essential media (DMEM) supplemented with 2% fetal bovine serum (FBS) and 1% penicillin/streptomycin. Cells were stored at 37°C, 5%CO2, and media was changed every other day.

### Overexpression and purification of FOLR3

Recombinant FOLR3 used in experiments was obtained by overexpression in HEK-293T cells and purification from the cell culture supernatant. HEK-293T cells were grown to 80% confluency in a 100mm cell culture dish and switched to DMEM media with 5% FBS. Cells were transfected with 5ug of FOLR3 expression plasmid with FLAG tag (Origene RC212963) using JetPRIME transfection reagent (Polyplus) following the manufacturer’s procedures. The cell culture media was collected and replaced every three days over two weeks. The collected media was then processed for FOLR3 purification. 30mL of media was concentrated to 3mL using 3K molecular weight cutoff filters (MWCO) from Millipore. FOLR3 was then purified using Pierce™ Anti-FLAG Magnetic Agarose following the manufacturer’s procedures. FOLR3 was eluted from magnetic agarose using 3X FLAG peptide and stored in PBS at -80°C. FOLR3 purity was assessed by gel electrophoresis, and protein concentration was determined using the Bradford assay.

### Western blot

LX-2 cells were grown in 6-well plates to 80% confluency. The cells were then serum-starved overnight and treated with FOLR3, TGFβ1, or vehicle for 24 hours. Cell culture supernatant was then collected and concentrated in 3K MWCO filters to ∼25ul. Samples were mixed with SDS sample buffer 4:1 and boiled at 95°C for 5 minutes. Samples were then separated on a 4-12% Bis-Tris gel and transferred to a nitrocellulose membrane at 100V for 40 minutes. The membrane was blocked with 5% blotting grade milk in Tris-buffered saline containing 0.05% Tween 20. They were then probed with primary antibodies for COL1A1 (Cell Signaling 39952S) 1:1,000. The membrane was washed with Tris-buffered saline 0.05% Tween 20 and probed with secondary antibody IR800DyeCW Goat anti-rabbit (LiCor) 1:5,000 for one hour. The membrane was then rewashed and imaged using LiCor Odyssey Imager.

### Immunofluorescence

Immunofluorescence experiments were performed in a 96-well plate. LX-2 cells were grown to 80% confluency and serum-starved overnight in DMEM. The cells were then stimulated with TGFβ1 5ng/mL for 2 hours. FOLR3 200nM was added, and after 30 minutes, HTRA1 200nM was added. The cells were then fixed in 4% paraformaldehyde for 10 minutes. After fixation, the cells were washed three times in PBS and blocked with 10% goat serum in PBS for one hour. Primary mouse monoclonal antibody for TGFBR2 (ProteinTech 66636-1-Ig) was diluted 1:100 in 1% bovine serum albumin and added to cells for two hours at 37°C. Cells were washed three times in PBS and incubated with polyclonal goat anti-mouse Alexa555 secondary antibody for 1 hour at 37°C. Cells were washed three times in PBS and imaged using Operetta CLS high content imager. Alexa555 intensity was measured in the membrane region of each cell.

### Immunoprecipitation

Immunoprecipitation of FOLR3 with a FLAG tag was performed to identify protein interactors with FOLR3. FOLR3 was added to PierceTM Anti-DYKDDDDK magnetic agarose (Invitrogen) to immobilize FOLR3. 200ug of HSC protein lysate was added to the agarose and incubated for one hour at 37°C. The beads were then washed with PBS three times. FOLR3 was eluted from the beads with 3X FLAG (ApexBio A6001). The eluates were then prepared for LC-MS/MS analysis described in the global proteomic analysis. The samples were eluted with LDS sample buffer for Western blot analysis and processed as described in the Western blotting section with primary antibodies for FLAG (R&D MAB8529) and HTRA1 (ProteinTech 55011-1-AP).

### Animal studies

Animal experiments were approved by the Institutional Animal Care and Use Committee (IACUC). Specific pathogen-free, 6-week-old, male, C57BL/6NTac mice of approximately 25 g body weight were purchased from Taconic Biosciences. The animals were weighed and ear-tagged seven days after arrival for individual identification. Animals were randomly assigned into three groups (n=3 per group), with no statistical difference in body weight. At seven weeks, the NASH group started receiving an Amylin diet consisting of 40 kcal% fat, 20 kcal% fructose, and 2% cholesterol (Research Diets # D09100310i). Control animals received a CHOW diet. At week 41, FOLR3 animals began receiving 1ug of recombinant FOLR3 once a day by i.p. injection. Animals were euthanized at 45 weeks, and liver tissue was stored at −80 °C.

### Liver Histology

Formalin-fixed liver tissues were paraffin-embedded and sectioned onto slides. Slides from each animal were stained with hematoxylin and eosin (H&E) and Sirius red staining. H&E and Sirius red staining quantitated the NAFLD activity score (NAS) and fibrosis score, respectively. The scores were quantitated based on the scale developed by the Nonalcoholic Steatohepatitis Clinical Research Network^57^. The parameters evaluated were steatosis, lobular inflammation, hepatic ballooning, and fibrosis.

### Statistical analysis

Data are expressed as mean values ± standard mean error (SEM). Statistical comparisons were performed using unpaired, two-tailed Student’s t-tests. Differences were considered significant when P<0.05.

