## Supplementary figures and images for "Secreted folate receptor-gamma drives fibrogenesis in nonalcoholic steatohepatitis by amplifying TGFβ signaling in hepatic stellate cells"

### Extended Data Figure 1

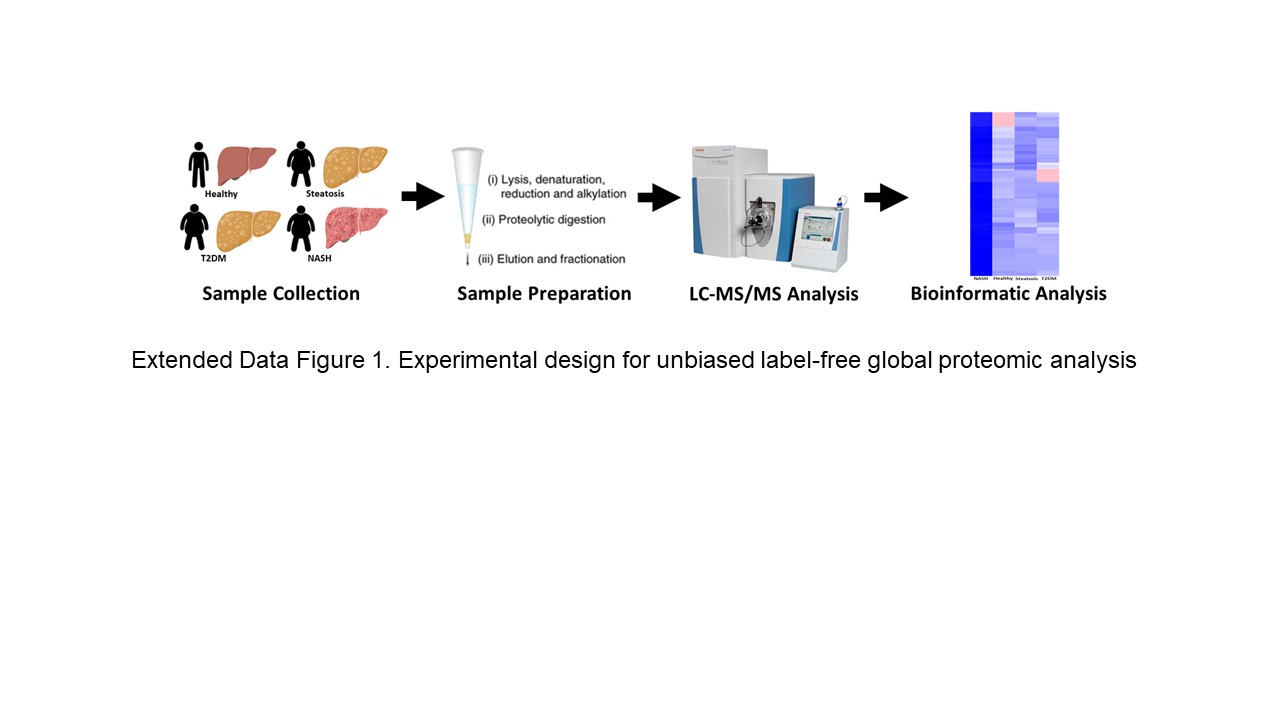

### Extended Data Figure 2

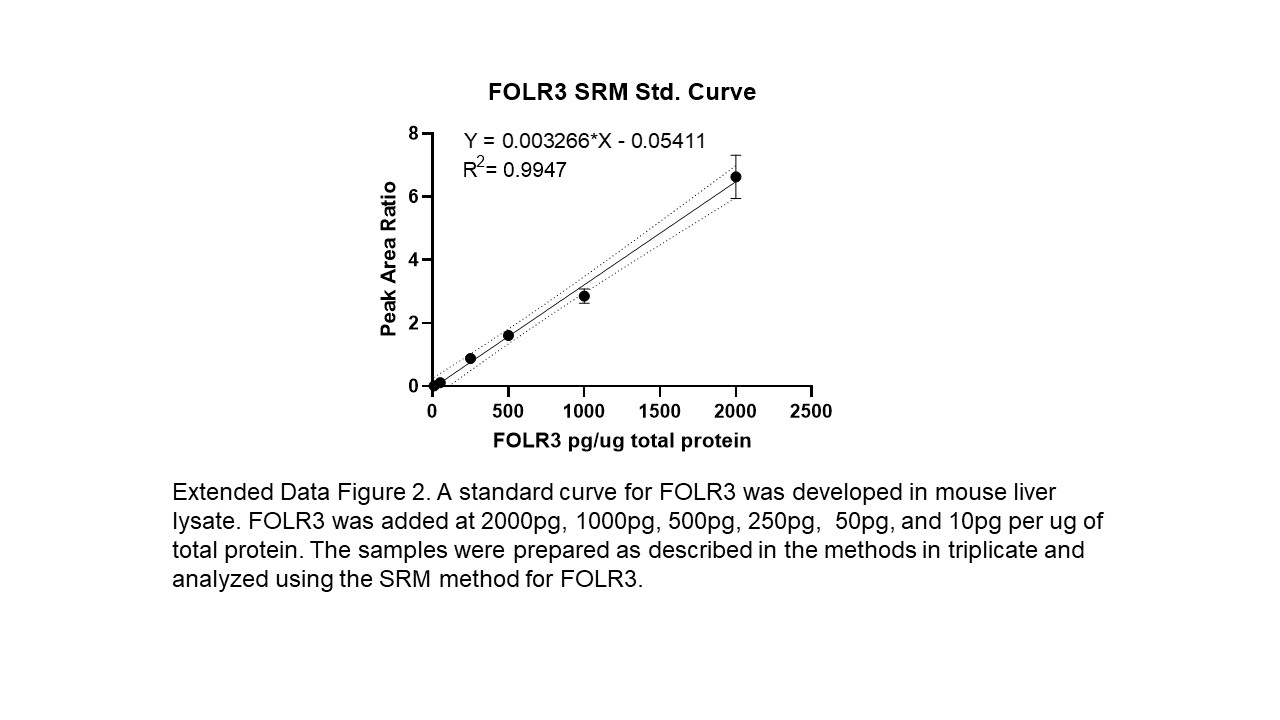

### Extended Data Figure 3

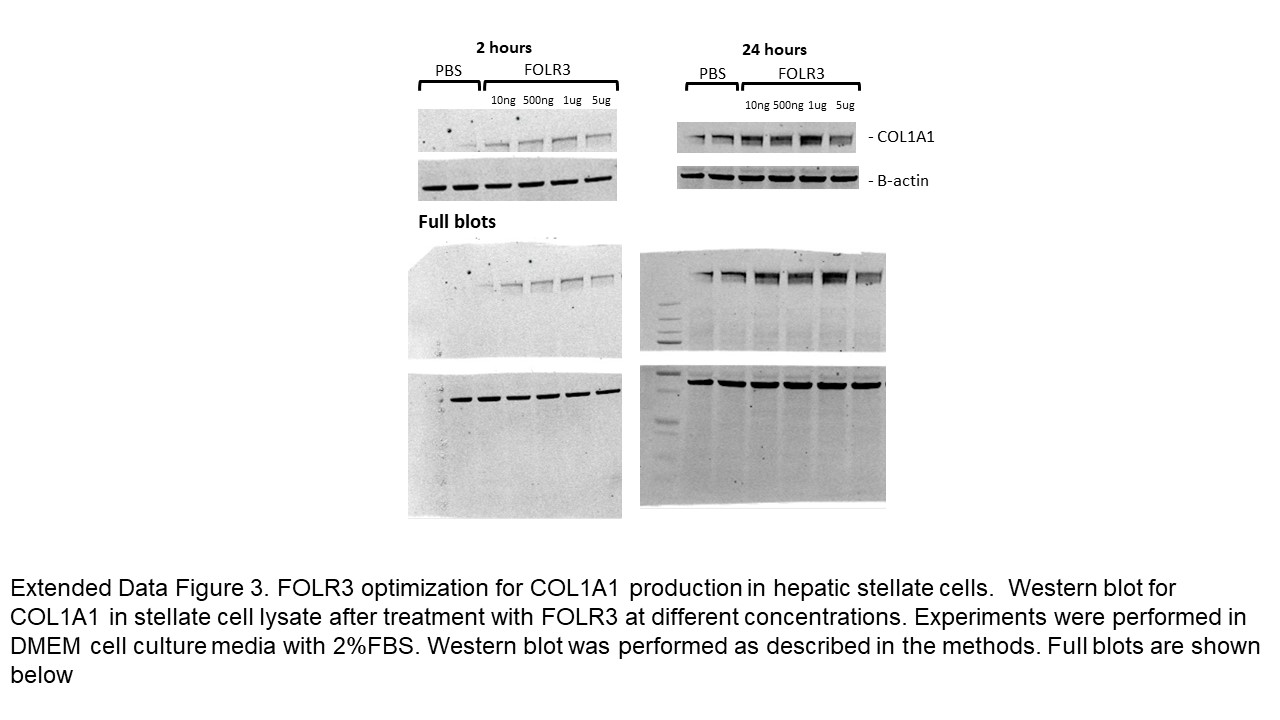

### Extended Data Table 3

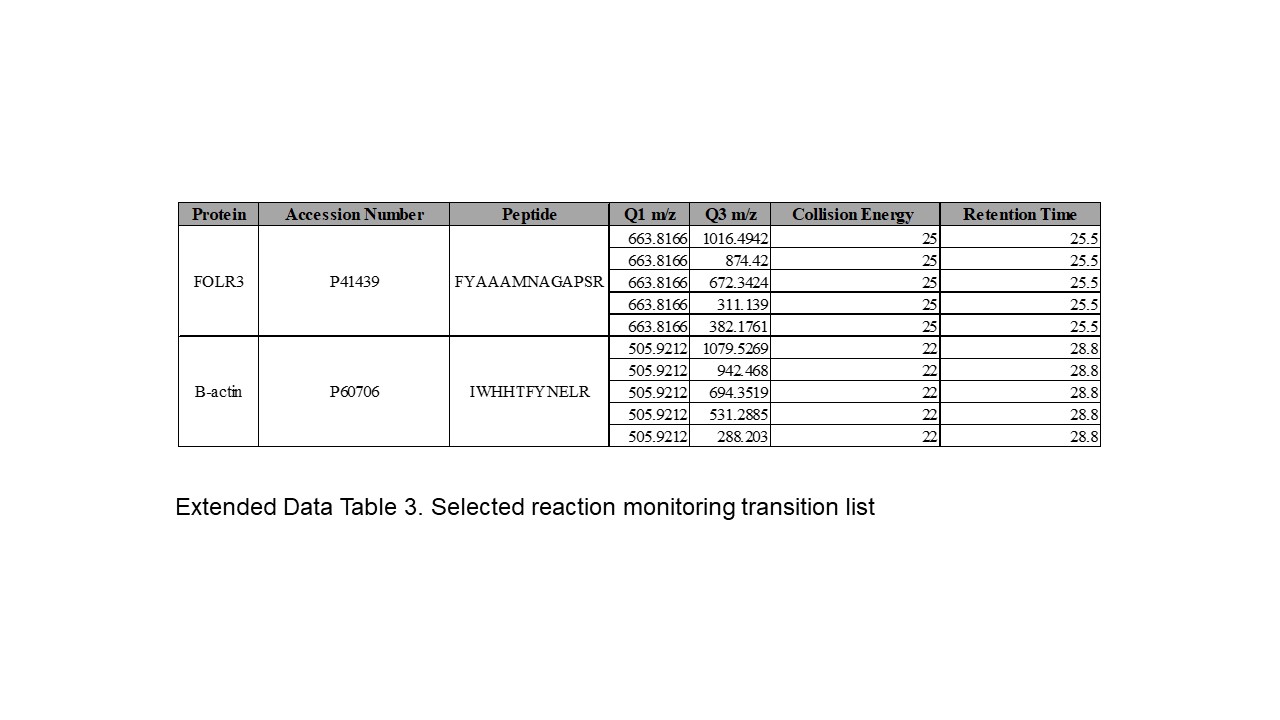

### Extended Data Table 5

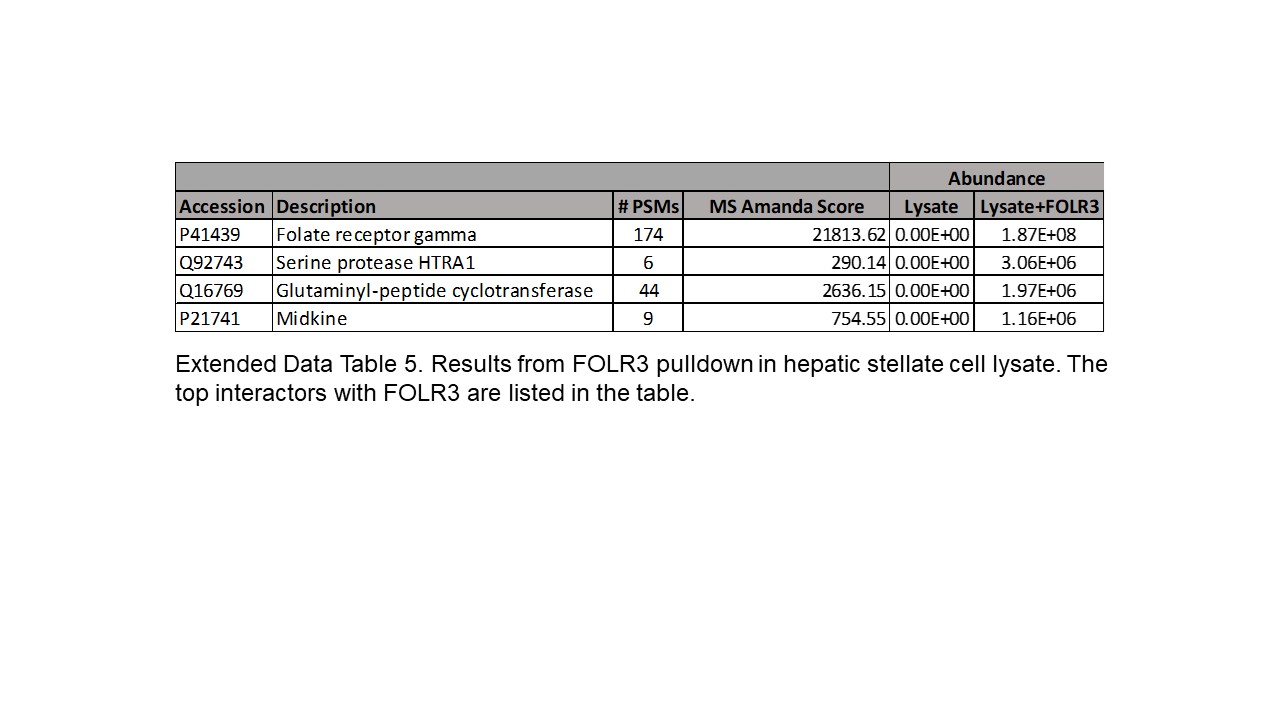
